# A cell type-specific multiomics uncovers a guard cell-specific RAF15-SnRK2.6/OST1 kinase cascade

**DOI:** 10.1101/2021.07.06.451220

**Authors:** Hongliang Wang, Yubei Wang, Rongxia Li, Weiwei Ren, Tian Sang, Bing Zhao, Xiao Wang, Xuebin Zhang, Shaojun Dai, Chuan-Chih Hsu, Chun-Peng Song, Pengcheng Wang

## Abstract

Multicellular organisms such as plants contain different cell types with specialized functions. Analyzing the characteristics of each cell type reveals specific cell functions and enhances understanding of organization and function at the organismal level. Here we report a highly-sensitive and efficient cell type-specific multiomics pipeline, combining simplified flow cytometry-based cell sorting for fluorescent protoplasts and optimized nanoscale proteomics and metabolomics methods, which allow in-depth analysis of the proteomes and metabolomes of a particular cell type. Using this method, we quantitatively compared the proteomes and metabolomes of guard cells and mesophyll cells and revealed that the enrichment of signal transduction-related proteins enables guard cells to respond rapidly to various environmental stimuli. We uncovered a guard cell-specific kinase cascade whereby RAF15 and OST1 mediate ABA-induced stomatal closure. This pipeline can be applied to various cell types in plant or non-plant systems to learn how cells function in highly organized multicellular organisms.

## INTRODUCTION

Multicellular organisms such as plants contain different cell types with specialized structures and functions. Dissecting the specific structural and functional information in each cell type is essential to understanding how a particular cell type works and how different cells integrate into a complex organism. Recently, the breakthrough of nucleic-acid-amplification-based DNA and RNA sequencing technologies allowed us to mine the transcriptional heterogeneity in different plant tissues ^1–3^. Generally, current single-cell transcriptome techniques use high-throughput and limited depth sequencing of thousands of cells, which enables rapid identification of rare and novel cell types, simultaneous characterization of multiple different cell types and states, and identification of unique marker genes in each cluster/cell type ^1^. After understanding the transcriptional heterogeneity of cells, it is necessary to perform in-depth analyses of the proteome, including post-translational modifications (PTMs), and metabolome to understand the specialized structure, function, signal transduction, and metabolism in a particular type/cluster of cells.

However, because proteins cannot be amplified *in vitro* like DNA/mRNA, even cutting-edge mass spectrometry can only identify about 1,000 proteins in a single mammalian cell ^4,5^. Comprehensively analyzing the proteome in a single cell is still challenging ^6^. Alternatively, enriching for a particular type of plant cell is a realizable approach for in-depth proteomics ^7^. However, until now, such single-cell-type proteomics was mainly performed in limited cell types that can easily be isolated, including pollen ^8–11^, guard cell (GC)^12–14^, and mesophyll cell (MC). GC is a specialized epidermal cell that plays an important role in gaseous exchange in and out of plant leaves by regulating the opening and closing of the stomas. Using GC protoplasts (GCP) isolated by enzyme digestion and centrifugation, several groups performed pioneering single-cell-type proteomic studies in Arabidopsis, *Brassica napus*, and maize, and discovered unique proteins, signal transduction pathways, or metabolism that determine the particular function and development of GC ^12–15^. Using fluorescence-activated cell sorting (FACS) and GeLC-MS/MS (in-gel tryptic digestion followed by liquid chromatography-tandem mass spectrometry), proteomic analyses identified nearly 2,000 proteins from six different cell types in the Arabidopsis root tips ^16^. A similar study compared the proteomes in epidermal and inner cell lines in Arabidopsis root ^17^. However, current methods are applicable to limited cell types and require millions of cells, and the efficiency of cell acquirement and the sensitivity of proteomics needs to be improved.

Here, we used a FACS method to enrich for fluorescent protoplasts. Combining with an optimized nanoscale label-free MS analysis method, we set up a cell type-specific pipeline applicable to proteomics and metabolomics studies of GC and MC, as well as root epidermal cells in Arabidopsis and rice. This pipeline allowed us to identify more than 4,500 proteins and 1,600 metabolites in guard cell protoplasts (GCP) and mesophyll cell protoplasts (MCP), using standard commercial mass-spectrometry machines. Quantitative comparison of GCP and MCP revealed the enrichment of signaling components in GC that may enable GC to respond to various environmental stimuli quickly. We uncovered a unique GC-specific RAF15-SnRK2.6 kinase cascade based on these data, which has a crucial role in ABA-induced stomatal closure. This pipeline can be applied to various cell types in plant or non-plant systems to learn how cells work in highly organized multicellular organisms.

## RESULTS

### A cell-sorting-based proteomics pipeline for limited plant samples

We set up a FACS method to enrich for fluorescent protoplasts (Figure 1A). We used the *pGC1::GFP* transgenic plants ^18^, which carry GC-specific expressed GFP, as a model system to test our pipeline. The *pGC1::GFP* leaves were digested into protoplasts, and after removal of tissue fragments by filtration, the protoplasts were subjected to FACS (Figures 1B and 1C). This simplified cell sorting procedure works well to isolate hundreds to millions of GCP from Arabidopsis leaves. We typically isolated up to 10,000 fluorescent GCP from 0.2 g leaf tissue (about 4-6 mature leaves from 3-week-old seedlings). We also optimized a nanoscale protein extraction and digestion procedure (Figure 1A). The enriched fluorescent GCP were lysed in sodium lauroyl sarcosinate (SLS)-sodium deoxycholate (SDC) buffer and directly subjected to trypsin digestion in a single microcentrifuge tube to avoid protein loss in precipitation and transferring (Figure 1A). After one-step desalting on C18 Stage-Tip, the peptides were subjected to LC-MS/MS analysis (Figure 1A). Compared to the guanidine chloride (Gdn-Cl) method optimized for medium-scale plant tissue extract ^19,20^, the simplified SLS-SDC method allowed us to identify more peptides/proteins, especially for hundreds to thousands of protoplasts. For example, we identified 2,827 peptides, representing 1,012 proteins, from about 500 GCP with the SLS-SDC method, which is nearly two-fold more than with the Gdn-Cl method (Figures 1D, 1E and S1, Dataset S1). The Gdn-Cl method acquired more peptides/proteins from 5,000, and 10,000 protoplasts (Figures 1D and S1), probably because precipitation and resuspension remove metabolites and improve the subsequent enzyme digestion and MS detection. Therefore, our pipeline is suitable for proteomics analysis in samples with limited material.

**Figure 1.**
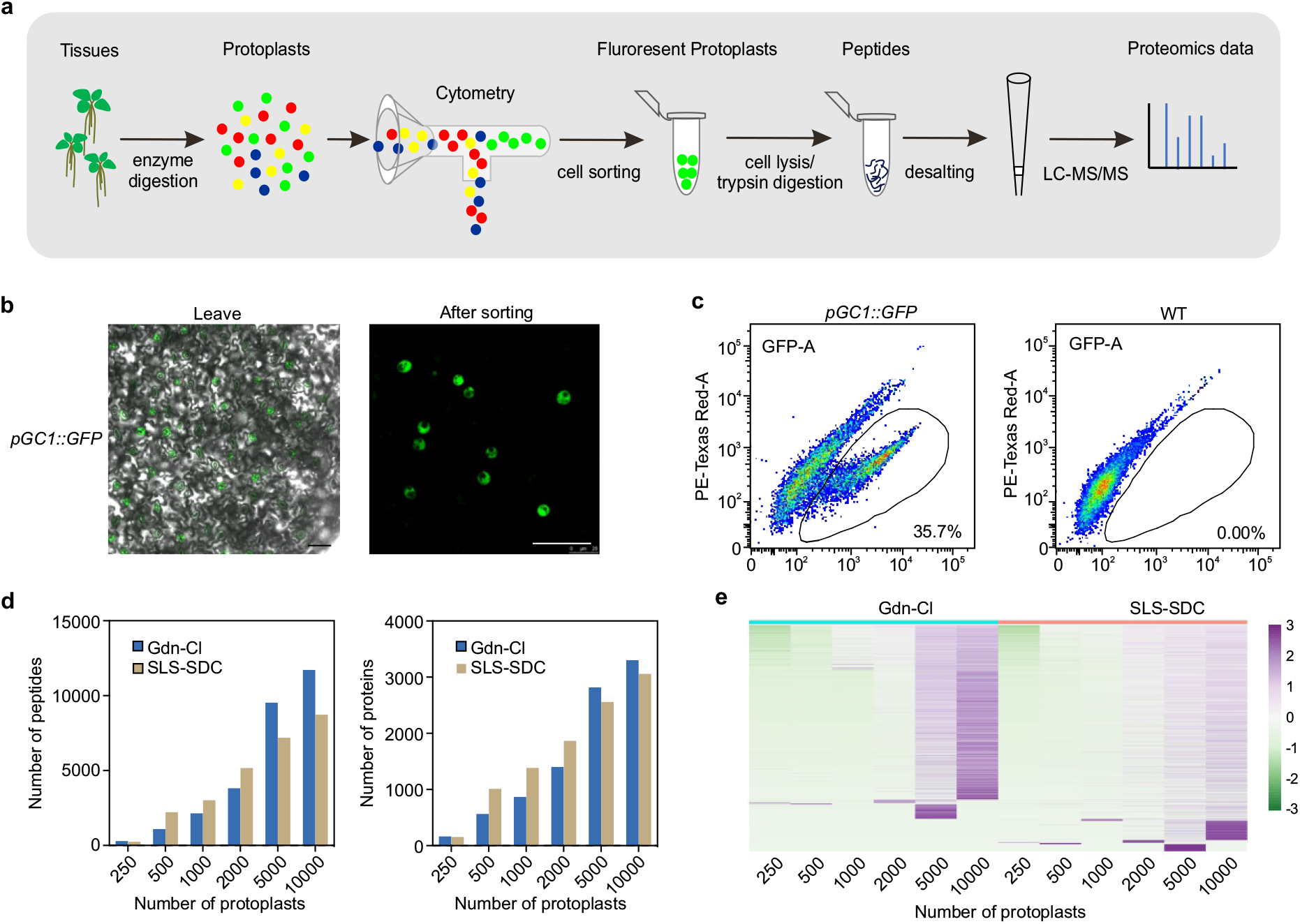
The cell-sorting-based nano-scale pipeline for isolating and proteomic analysis of guard cell protoplasts. **a.** Workflow of the cell-sorting-based nanoscale pipeline for single-cell (type) proteomics. The fluorescent protoplasts were sorted by cytometry after enzymatic digestion and filtration. Protein extraction and trypsin digestion were performed in a single microcentrifuge tube, and after desalting through the C18 Stage-Tip, the peptides were subjected to LC-MS/MS. **b.** Fluorescent images showing the GFP fluorescence of *pGC1::GFP* transgenic leaves (left, Bar=50 μm) and sorted GCPs (right, Bar=50 μm). **c.** Representative flow cytometry scatter plots for *pGC1::GFP* (left) and wild type control (WT, right). **d.** The number of peptides (left) and proteins (right) identified from the indicated number of GCPs. **f.** Heatmap showing the number of peptides in each protein detected by SLS-SDC or Gdn-Cl buffers from the indicated number of GCPs.

### The proteomics pipeline can be applied to different cell types

To test if this pipeline is applicable to other types of fluorescent cells, we enriched for protoplasts of root cells in *DR5 rev::GFP* transgenic Arabidopsis (Figures 2A and 2B), a marker line for auxin response ^21^. We obtained an average of 6,871 peptides, representing 1,950 proteins, from approximately 1,000 fluorescent protoplasts of *DR5 rev::GFP* (Figure 2C and Dataset S2). GO analysis revealed that proteins involved in root development, auxin transport, translation, response to cold, oxidative stress, and embryo development are enriched in *DR5 rev::GFP* fluorescent protoplasts (Figure S2A and Dataset S2).

**Figure 2.**
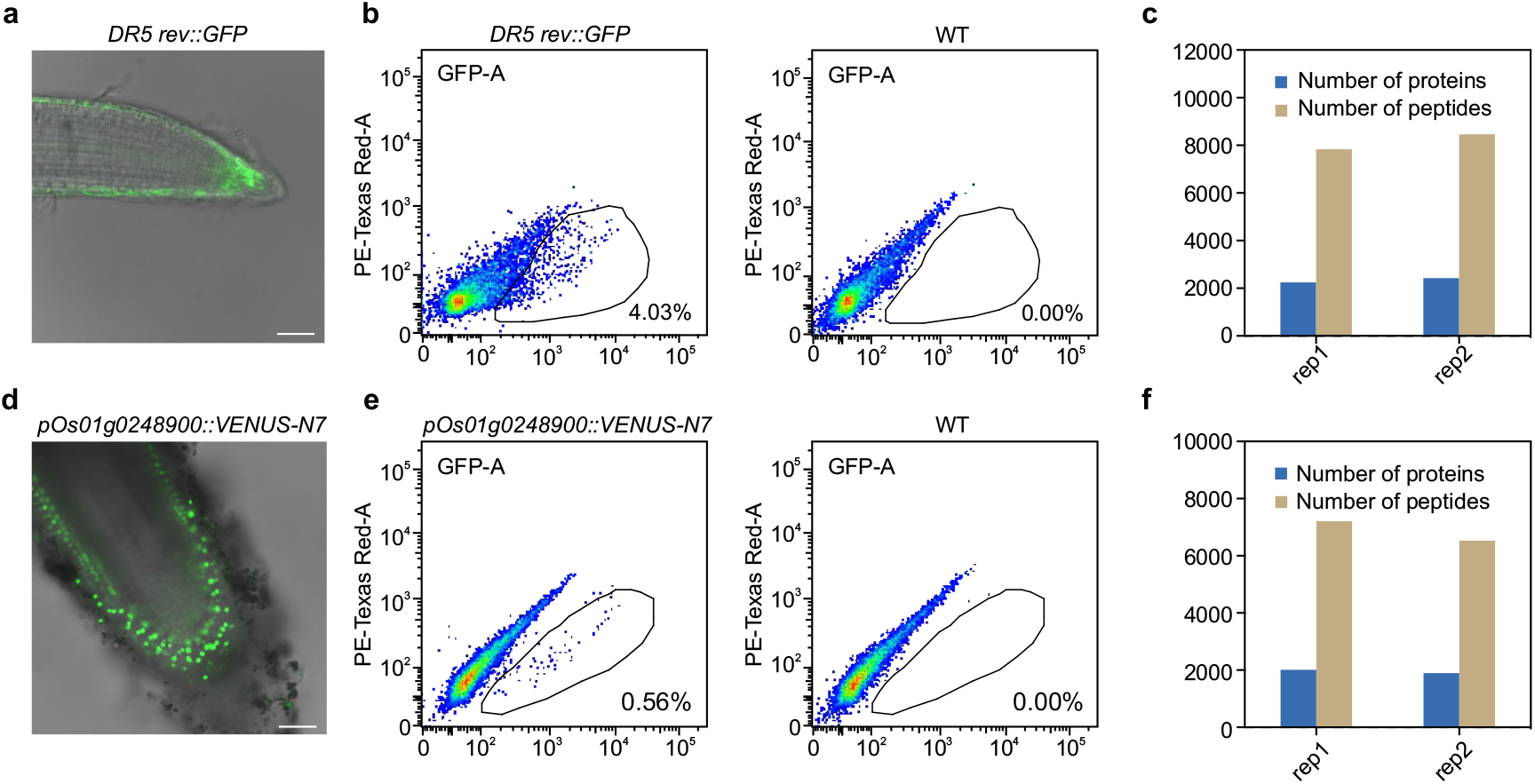
Proteomic analysis of *DR5 rev::GFP* and *pOs01g0248900::VENUS-N7* fluorescent protoplasts. **a.** Image showing the fluorescence of *DR5 rev::GFP* transgenic seedling roots. Bar=50 μm. **b.** Representative flow cytometry scatter plots for *DR5 rev::GFP* (left) and wild type control (right). **c.** The number of peptides and proteins identified from approximately 1,000 *DR5-rev::GFP* fluorescent protoplasts. **d.** Image showing the fluorescence of root in *pOs01g0248900:: VENUS-N7* transgenic rice seedling. Bar=50 μm. **e.** Representative flow cytometry scatter plots for *pOs01g0248900:: VENUS-N7* (left) and wild type (WT, right). **f.** The number of peptides and proteins identified from approximately 1,000 *pOs01g0248900:: VENUS-N7* transgenic rice fluorescent protoplasts.

A recent single-cell transcriptome study in rice revealed that *pOs01g0248900:: VENUS-N7* is strongly expressed in atrichoblasts in rice root tips ^22^. We obtained 8,149 peptides, representing 2,232 proteins, from approximately 1,000 fluorescent protoplasts of *pOs01g0248900::VENUS-N7* (Figures 2D-F, Dataset S3). Consistent with the finding that some atrichoblast-enriched genes are involved in cell wall biosynthesis ^22^, we found server GO terms like Go:0071669 (plant-type cell wall organization or biogenesis) and GO:0071554 (cell wall organization or biogenesis) in the proteins identified in the fluorescent protoplasts of *pOs01g0248900::VENUS-N7* (Dataset S3). Other enriched GO terms included oxoacid and cellular ketone metabolic processes, transport, and glycolysis (Figure S2B and Dataset S3). Therefore, our pipeline is useful for studying a variety of plant cell types.

### In-depth proteomics analysis of GCP and MCP

Encouraged by this result, we further performed label-free quantitative proteomics from 1 μg total protein extract from GCP (about 30,000 cells) and MCP (about 15,000 cells) to acquire in-depth proteomic profiles of GCP and MCP. We identified a total of 27,511 peptides, representing 4,622 proteins, in GCP, and 25,235 peptides, representing 4,237 proteins, in MCP (Figure 3A, Dataset S4 and S5). Among these proteins, 526 proteins were identified at least once in the three biological replicates of GCP, but not in any sample of the MCP, and 141 proteins were only detected in MCP (Figure 3A, Dataset S4 and S5). We also performed a quantitative comparison of the GCP and MCP proteomes (Dataset S6-8). As shown in Figures 3B and 3C, 2,483 proteins showed enrichment in GCP (FC > 2, *p* < 0.05) (Dataset S7). Some proteins involved in stomatal development/function were present in the list of GCP enriched proteins (Figures 3D-F). FAMA and Inducer of CBF Expression 1 (ICE1)/ SCREAM (SCRM) govern GC mother division and promote GC differentiation ^23^. FAMA, ICE1/SCRM, and their interacting protein, the Third Largest Subunit of Nuclear DNA-dependent Pol II (NRPB3), were present in the list of GCP enriched proteins (p < 0.05) ^24^. After forming GC, STOMATAL CARPENTER 1 (SCAP1) is required to develop functional stomata ^25^. MYB DOMAIN PROTEIN 60 (MYB60) involved in the stomatal movement is a known target of SCAP1 ^25^. Both SCAP1 and MYB60 were only present in GCP, not in MCP, in our study. Besides these proteins for stomatal development, we also noticed that known regulators of GC intensity, like MUS, SCD1, and PHYB, were enriched in GC (Figure 3E). Many key components of stomatal movement regulation in response to blue light (BL), high CO_2_, and abscisic acid (ABA) were also enriched in GCP (Figure 3F). For example, BETA CARBONIC ANHYDRASE 4 (BCA4) ^26^, HIGH LEAF TEMPERATURE 1 (HT1) ^27^, and CONVERGENCE OF BLUE LIGHT AND CO2 1 (CBC1)/CBC2 ^28^, mediating CO_2_-induced stomatal movement, and blue light receptors CRYPTOCHROME 2 (CRY2) and PHOT2 ^29^, the PHOT2 substrate BLUE LIGHT SIGNALING1 (BLUS1) ^30^ and BLUE LIGHT-DEPENDENT H^+^-ATPase PHOSPHORYLATION (BHP) ^31^, were present in the list of GCP enriched proteins. A recent study suggested that glucose, not malate, is the major starch-derived metabolite in GC for rapid stomatal opening ^32^. The b-AMYLASE1 (BAM1) ^32^, a glucan hydrolase, and ATP-BINDING CASSETTE B14 (ABCB14) ^33^, the malate transporter, were enriched in GCP. Consistently, the enriched KEGG pathway and enriched GO terms in the GCP were mainly related to transport, RNA degradation, and responses to ABA, cold, and osmotic stress (Figures 3G, S3, and Dataset S9).

**Figure 3.**
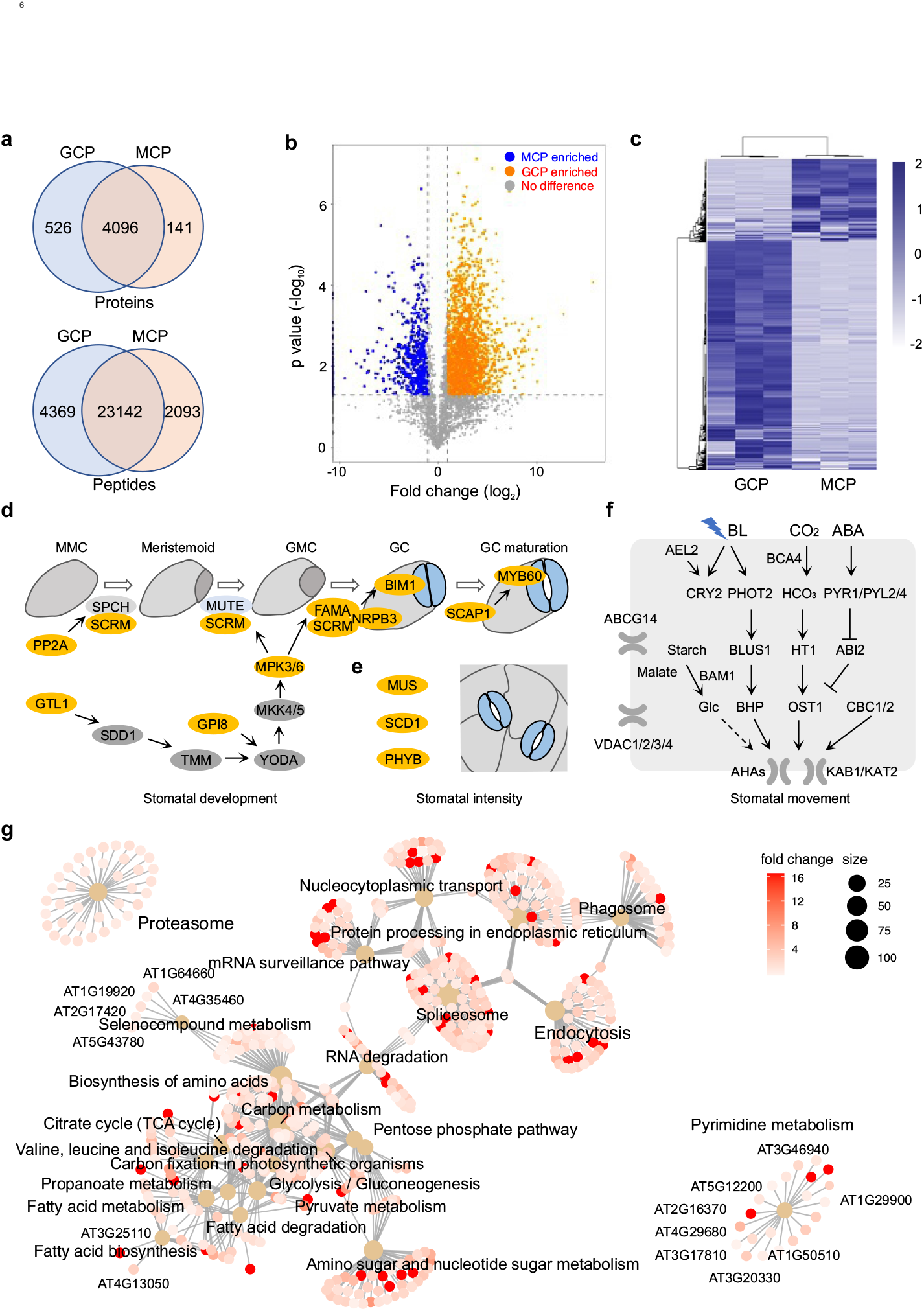
Quantitative comparison of GCP and MCP proteome. **a.** Venn diagram showing the overlap between the proteins and peptides identified from GCP and MCP proteomes. **b.** Volcano plot of differentially up-regulated (upper panel) or down-regulated (bottom panel) proteins in the proteome of GCPs and MCPs. On the y-axis, the negative log10 values are plotted. On the x-axis, the log2 values of the fold change are seen in the comparison. **c.** Heatmap showing the quantitative comparison of proteins in GCP and MCP proteomes. **d**. Some known proteins that function in GC development are present in the proteome of GCP. **e**. Some known proteins that function in the regulation of GC intensity are present in the proteome of GCP. Yellow circles highlight proteins detected in GCPs. **f.** Some known proteins that function in stomatal movement in response to ABA, CO_2_ and blue light (BL) are enriched in GCPs. All proteins listed in this panel show higher abundance in GCPs than that in MCPs (FC > 2, *p* < 0.05). **g.** Cnetplot generated by ClusterProfiler indicating proteins associated with enriched KEGG pathways; dots in light brown represent KEGG pathways, with the size of dots determined by the number of proteins belonging to the pathway. Small dots of the same size represent proteins, the dot color is determined by fold changes of protein abundance of GCP/MCP.

There were 664 proteins only present or enriched (FC > 2, *p* < 0.05) in MCP (Figures 3B, 3C, and Dataset S8). Notably, 546 out of these 664 proteins are known to be components of chloroplast (GO term 0009507). Other enriched GO terms of MCP enriched proteins included photosynthesis, photorespiration, pigment, chlorophyll metabolic process, electron transport chain, and response to radiation and light stimulus. These data supported the dominant role of MC in photosynthesis (Dataset S9).

### Metabolomics analysis of GCP and MCP

To determine the metabolite differences between GCP and MCP, we also performed untargeted metabolomics using approximately 100,000 GCP or MCP and detected 1,678 metabolites (Dataset S10). Principal component analysis indicated a clear separation of the metabolomes in the two different cell types (Figure 4A). We found 260 metabolites significantly (log_2_ ratio > 0.26, p < 0.05, unpaired two-tail *t*-test) enriched in GCP compared to MCP, and 530 metabolites enriched in MCP compared to GCP (Figures 4B and 4C, Dataset S10). Pathway enrichment analysis of the differentially accumulated metabolites indicated an over-representation of metabolites derived from flavone and flavonol biosynthesis (Figure 4D, Dataset S10).

**Figure 4.**
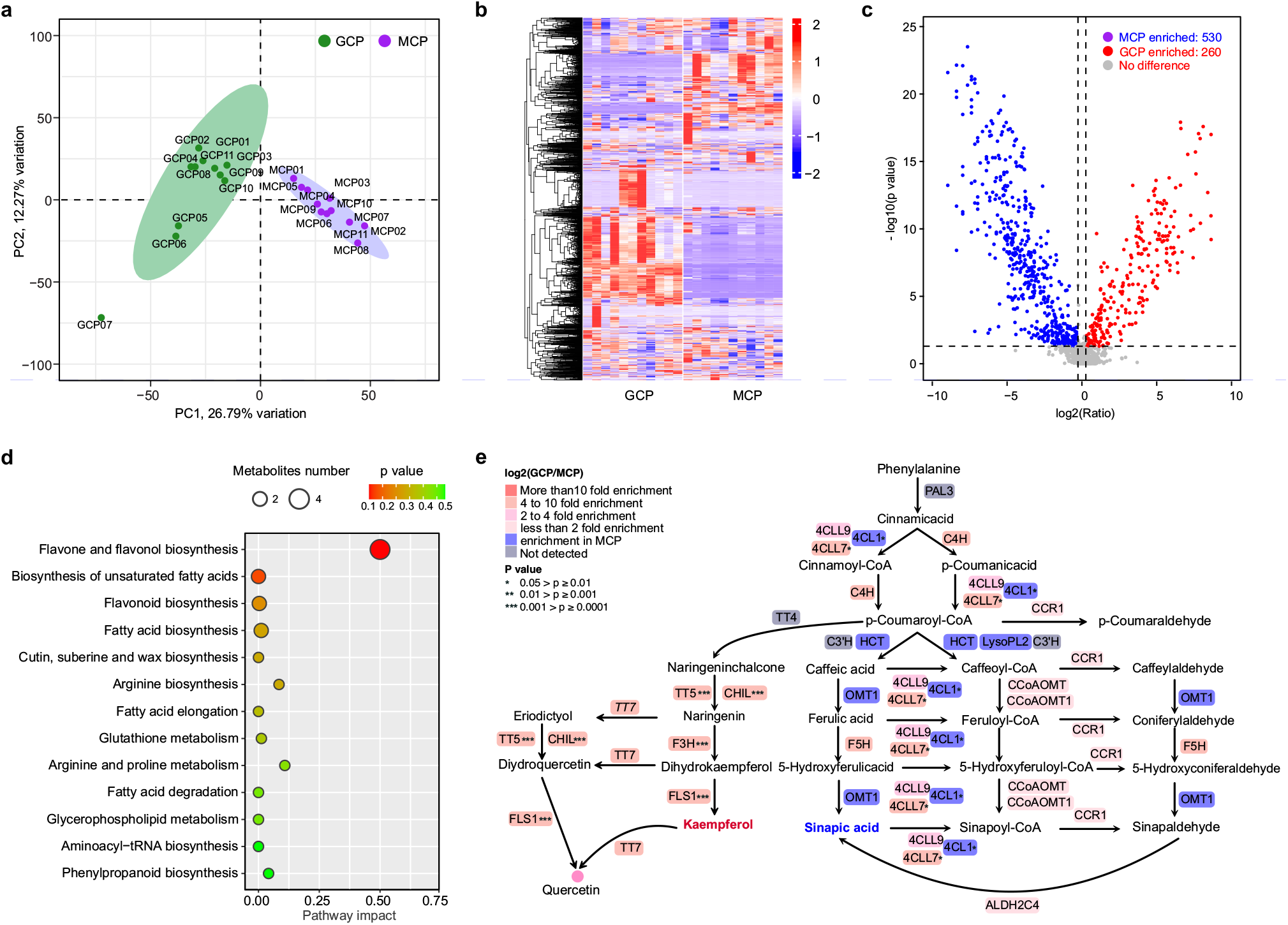
Quantitative comparison of GCP and MCP metabolome. **a.** PCA analysis of metabolome of the 11 MCP and 11 GCP samples. **b.** Heatmap showing the quantitative comparison of metabolites in GCP and MCP metabolome. **c.** Volcano plot showing the quantitative comparison of metabolites in GCP and MCP metabolome. **d.** Pathway analysis of metabolites differentially accumulated in GCP and MCP. **e.** The protein-metabolite joint pathway analysis of phenylalanine/flavonoid metabolism between GCP and MCP. The metabolites, proteins, and their relationships are derived from the phenylpropanoid biosynthesis pathway (ath:00940) and the flavonoid biosynthesis pathway (ath:00941) in the KEGG database. The enzymes enriched in GCP or MCP, respectively, are colored with red or blue. Grey indicates that the enzyme has not been detected by proteomics, but it is indispensable in phenylpropane metabolism pathway.

We also performed an integrative analysis combining the proteomics and metabolomics data to reveal the GC-specific metabolic pathway. Several intermediates in flavonoid/phenylpropanoid metabolic pathways, like Kaempferol-3-glucoside-7-O-rhamnoside, Kaempferol, Rutin, and Afzelin, were highly enriched in the GCP (*p* < 0.05, unpaired two-tailed *t*-test) (Figure S4A). Consistently, the enzymes catalyzing flavonoid/phenylpropanoid biosynthesis, like Flavonol Synthase 1 (FLS1), Flavanone 3-hydroxylase (F3H), Transparent Testa 5 (TT5), and Chalcone Isomerase Like (CHIL), were highly accumulated in GCP (Figure 4E and S4B). Previous studies suggested a crucial role of flavonoids in scavenging reactive oxygen species (ROS) in GC ^34–36^. The enrichments of these enzymes and metabolites suggest their potential roles in regulating flavonoid metabolism and ROS homeostasis in the GC.

The quick opening and closing of stomata largely depend on the structure of the cell walls ^37^. In *gpat4/gpat8*, a mutant lacking two glycerol-3-phosphate acyltransferases (GPAT) required for cutin biosynthesis, the cuticular ledges between GC are defect ^38^. Our data showed that several enzymes in cutin biosynthesis, including GPAT4, GPAT8, and a short-chain dehydrogenase ECERIFERUM 3 (CER3), are highly accumulated in GCP (Dataset S7). Supporting this notion, some enzymes and metabolites in the fatty acid biosynthesis and elongation pathway were highly accumulated in GCP (Figure 4D and S4B).

### GC-enriched RAF15 mediates OST1 activation and stomatal closure

We then systemically surveyed the presence of known ABA signaling components in GCP and MCP. Dozens of known ABA signaling components showed a GC-enriched pattern, and only very few components had a higher abundance in MCP than in GCP (Figure 5A). For example, the ABA receptors PYR1, PYL1, and PYL2 showed 65.8-, 5.6-, and 446-fold enrichment, respectively, in GCP, while PYL4 was only detected in the GCP, not in the MCP (Figure 5B). SnRK2.6/OST1 (a master regulator of stomatal movement) and ABI2 showed 34.0- and 4.9-fold enrichment, respectively, in GCP (Figure 5B). By contrast, ABI1 showed an accumulation in MCP (Figure 5B). To our surprise, the B2 and B3 subgroup RAF kinases, known to phosphorylate and activate SnRK2.2, SnRK2.3, and SnRK2.6/OST1 in response to ABA ^39–41^, were not detected/enriched in our proteomics result (Figure 5B). This suggested that kinases other than B2 and B3 RAFs may contribute to ABA-induced SnRK2.6/OST1 activation and stomata closure.

**Figure 5.**
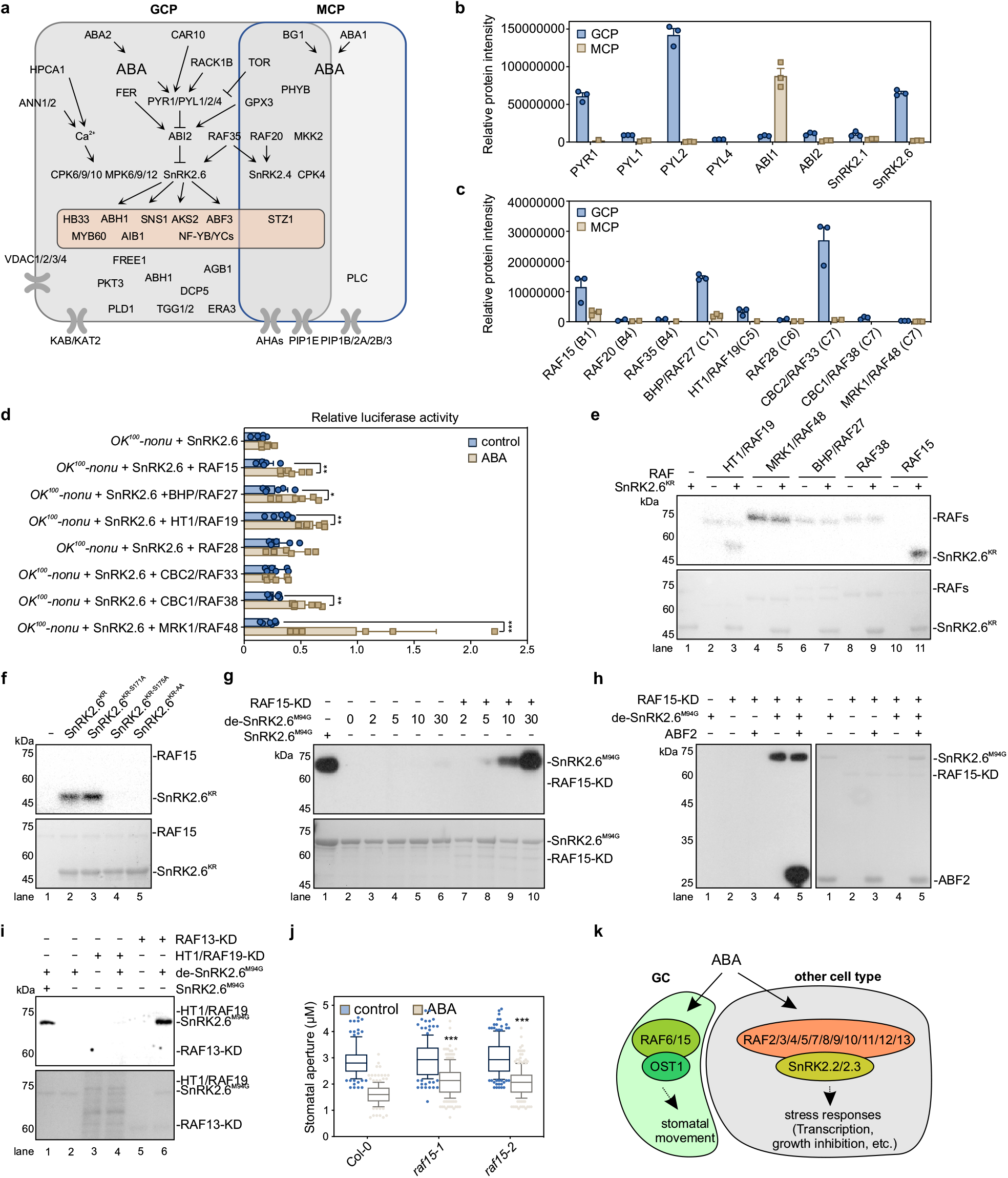
RAF15 mediated SnRK2.6/OST1 activation and ABA signaling in GC. **a.** Presence of ABA signaling components in the GC and MC proteomes. **b.** Relative protein abundance of ABA receptor PYR1/PYLs, PP2C phosphatases, and SnRK2s in the GC and MC proteomes. **c.** Relative protein abundance of B and C subgroup RAFs in the GC and MC proteomes. **d.** Activation of the reporter gene by the combinations of RAFs with SnRK2.6 in the protoplasts of *OK^100^-nonu*. The ratio of *RD29B-LUC* expression in the protoplasts with 5 μM ABA relative to that without ABA treatment was used to indicate the activation activity of different RAFs on SnRK2.6. Error bars, SEM (n = 6 individual transfections). **e.** Recombinant full-length RAF was used to phosphorylate SnRK2.6^KR^ (SnRK2.6^K50R^, a kinase-dead form of SnRK2.6), in the presence of [γ-^32^p]ATP. Autoradiograph (upper) and Coomassie staining (bottom) show phosphorylation and loading, respectively, of purified GST-HT1/RAF19, GST-MRK1/RAF48, GST-BHP/RAF27, GST-RAF38, HIS-RAF15, and HIS-SnRK2.6^KR^. **f.** Ser175Ala mutation abolished RAF15 phosphorylation on SnRK2.6^KR^. Recombinant full-length HIS-RAF15 was used to phosphorylate HIS-SnRK2.6^KR^, SnRK2.6^KR^ with Ser171Ala mutation (SnRK2.6^KR-S171A^), SnRK2.6^KR^ with Ser175Ala mutation (SnRK2.6^KR-S175A^), or SnRK2.6^KR^ proteins with Ser171AlaSer175Ala mutations (SnRK2.6^KR-AA^), in the presence of [γ-^32^p]ATP. Autoradiograph (upper) and Coomassie staining (bottom) show phosphorylation and loading, respectively, of purified RAF and HIS-SnRK2.6^KR^. **g.** RAF15-KD triggers the autophosphorylation of pre-dephosphorylated GST-SnRK2.6^M94G^ (de-SnRK2.6^M94G^). Anti γ-S immunoblot (upper) and Coomassie staining (lower) show thiophosphorylation and loading, respectively, of recombinant GST-RAF15-KD and GST-SnRK2.6^M94G^. **h.** RAF15-KD activates pre-dephosphorylated GST-SnRK2.6^M94G^ (de-SnRK2.6^M94G^) and the reactivated SnRK2.6^M94G^ phosphorylates itself and ABF2. Anti γ-S immunoblot (left) and Coomassie staining (right) show thiophosphorylation and loading, respectively, of recombinant GST-RAF15-KD, GST-SnRK2.6^M94G^, and GST-ABF2. Images shown are representative of at least two independent experiments. **i.** RAF13-KD, but not HT1, triggers the autophosphorylation of pre-dephosphorylated GST-SnRK2.6^M94G^ (de-SnRK2.6^M94G^). Anti γ-S immunoblot (upper) and Coomassie staining (lower) show thiophosphorylation and loading, respectively, of recombinant GST-RAF13-KD, GST-HT1/RAF19-KD and GST-SnRK2.6^M94G^. **j.** Stomatal closure of 4-week-old wild type and *raf15* mutants in response to ABA. Error bars, SD (n = 176 to 210 individual stomates). Unpaired two-tailed *t*-tests, *** p < 0.001. Box and whisker: 10–90 percentiles. **k.** Model showing the different RAFs mediate ABA signaling by phosphorylating SnRK2s in different cell type.

Our GCP proteomics identified RAF15 (AT3G58640) in the B1 subgroup, BHP/RAF27 in the C1 subgroup, HT1/RAF19 in the C5 subgroup, RAF28 in the C6 subgroup, and CBC1/RAF38, CBC2/RAF33, and AtMRK1/RAF48 in the C7 group (Figure 5C). Consistently, most of these genes showed higher expression in GCP than in MCP (Figures S5A and 5B). Using a transient expression assay system in MCP of *OK^100^-nonu*, which lacks most members of the B2 and B3 RAFs involved in ABA signaling, and has very low expression of SnRK2.6/OST1, we evaluated the function of these B1 and C subgroup RAFs in ABA signaling ^41^. Co-transfection of SnRK2.6/OST1 alone did not rescue the expression of ABA-induced *RD29B-LUC* in MCP of *OK^100^-nonu* (Figure 5D). However, co-transfection of RAF15, BHP/RAF27, HT1/RAF19, CBC1/RAF38, or MRK1/RAF48, but not RAF28 or CBC2/RAF33, rescued the expression of ABA-induced *RD29B-LUC* in MCP of *OK^100^-nonu* (Figure 5D). An *in vitro* kinase assay showed that full-length HT1/RAF19 or RAF15, out of the five tested RAFs, could directly phosphorylate SnRK2.6^KR^, a kinase “dead” form of SnRK2.6/OST1 (Figure 5E). Two serine residues in the activation loop of SnRK2.6/OST1, Ser171 and Ser175, are known to be phosphorylated by B2 and B3 RAFs, and their phosphorylation is essential for SnRK2.6/OST1 activation ^22,42,43^. Site-direct mutagenesis of Ser171 and Ser175 showed that RAF15 mainly phosphorylates the Ser175 in SnRK2.6 and the Ser175Ala mutation abolished the phosphorylation of SnRK2.6/OST1 by RAF15 (Figure 5F). Neither Ser171 nor Ser175 was the HT1/RAF19 phosphosite in SnRK2.6/OST1 (Figure S5C).

We then tested if RAF15 could reactivate the dephosphorylated inactive form SnRK2.6 (de-SnRK2.6). We used an adenosine triphosphate (ATP) analog-based *in vitro* kinase assay system, which can distinguish the trans- and autophosphorylation of SnRK2.6, to monitor the activity of SnRK2.6/OST1 ^41^. A Met94Gly (M94G) mutation in SnRK2.6/OST1 enlarges its ATP binding pocket, which can use the ATP analog N^6^-Benzyl-ATPγS to thiophosphorylate itself or its substrate ^41^. By this method, thiophosphorylation by activated SnRK2.6^M94G^ can be detected with an anti-thiophosphate ester antibody that only recognizes the thiophosphorylation. As shown in Figure 5G, pre-dephosphorylated SnRK2.6^M94G^ (de-SnRK2.6^M94G^) had no auto-thiophosphorylation activity (lanes 2-6). Application of the recombinant kinase domain of RAF15 (RAF15-KD) quickly induced the auto-thiophosphorylation of SnRK2.6^M94G^ in a time-dependent manner (lanes 7-10), suggesting that RAF15-mediated phosphorylation is sufficient to reactivate SnRK2.6^M94G^ *in vitro*, in the context of the auto-thiophosphorylation activity of SnRK2.6^M94G^. We then examined SnRK2.6^M94G^ activity by detecting the thiophosphorylation of ABA responsive element binding factor 2 (ABF2), a well-studied SnRK2 substrate. Adding RAF15-KD initiated the thiophosphorylation of SnRK2.6^M94G^ and ABF2 (Figure 5H, lanes 4 and 5). Thus, transphosphorylation by RAF15-KD is sufficient to reactivate the dephosphorylated SnRK2.6/OST1. Unlike RAF15-KD, HT1/RAF19-KD did not reactivate the dephosphorylated inactive de-SnRK2.6 in the *in vitro* activation system (Figure 5I, lane 3 and 4).

We also tested the ABA-induced stomatal closure in *raf15-1* and *raf15-2*, two T-DNA mutant lines of *RAF15* (Figure S5D). Both *raf15-1* and *raf15-2* showed a significantly impaired stomatal closure in response to ABA, compared to wild type (unpaired two-tailed *t*-test, p < 0.001) (Figure 5J). RAF13 in the B1 subgroup also phosphorylated Ser171 and reactivated SnRK2.6^KR^ *in vitro* (Figures 5I, lane 5 and 6 and S5E). Another B1 RAF, RAF14, had weak kinase activity and did not phosphorylate SnRK2.6^KR^ *in vitro* (Figure S5F). However, the abundance of *RAF13* and *RAF14* mRNA in GC was much less than that of *RAF15* mRNA (Figure S5A), and RAF13 and RAF14 protein was not detected in our GC proteomics (Dataset S4). We also noticed that all three members of the B1 RAF subgroup failed to phosphorylate SnRK2.4^KR^, one of the ABA-independent SnRK2s (Figure S5G). Thus, GC-enriched RAF15 may have a unique role in stomatal closure through specifically mediating ABA-induced SnRK2.6/OST1 activation (Figure 5K).

## DISCUSSION

Here, by optimizing the sample preparation and detection procedures, we developed a cell-sorting-based nanoscale pipeline that allows in-depth profiling of the proteome and metabolome of various cell types in plants. We can routinely isolate up to 10,000 protoplasts from a few leaves of Arabidopsis seedlings and identify more than 4,500 unique proteins and 1,600 metabolites in GCP. Compared to pioneering studies that identified approximately 2,000 proteins from 1,000,000 root cell protoplasts ^16^, and 2,000 proteins and 300 metabolites from 3 × 10^8^ GCP ^13,44^, our method showed a significantly improved sensitivity and efficiency. Importantly, this pipeline is also applicable to root cells in both Arabidopsis and rice, suggesting the potential of this pipeline in multiomics analysis in different cells from different plant species and even non-plant systems. Combined with the single-cell-transcriptomics that identify unique marker genes for each cell group, we may generate fluorescent transgenic lines and perform cell-specific multiomics in each cell type in a tissue or even organism. In the future, this pipeline may also be applied to the sorting and enrichment of different organelles for spatial proteomics to understand the localization and dynamics of proteins in cells ^45^. Such enriched cells or organelles could also be used for PTM proteomics to further reveal functional regulation at the proteome level.

As a FACS method, the protoplast isolation method for each cell type needs to be optimized. A few hours’ enzymatic digestion with cellulase/macrozyme and high-concentration mannitol may affect the stability or modification of some proteins, which limits the application of this pipeline in stimulus-responsive proteomics analysis. It is also limited to species than can be transformed. We noticed that some RAFs, e.g., RAF6, that are highly expressed in GC (Figure S5A) were not identified in our GC proteomics, and this might be due to degradation in protoplast/sample preparation. In addition, we failed to obtain enough quiescent center (QC) cells from *WOX5::ERGFP* transgenic plants, which might be due to the limitation of cytometry to isolate a cell type with such low abundance. In the future, new methods like magnetic-activated cell sorting may be helpful to isolate and perform multiomics analysis in such cell types.

The in-depth analysis of proteome and metabolome in GC allows us for the first time to uncover the molecular basis that determines the highly specialized structure and function of GC. SnRK2.6/OST1 is the master regulator in stomatal movement, and dysfunction of OST1 in *ost1* results in an open stomata phenotype even upon drought conditions or pathogen attacks ^46,47^. Under unstressed conditions, OST1 is dephosphorylated and inhibited by the A Clade PP2C phosphatases ^48^. Upon stress conditions, ABA binds to ABA receptor PYR1/PYL/RCAR proteins, and the ABA/receptor complex inhibits PP2Cs ^49–52^. Recombinant SnRK2.6 was considered to be auto-reactivated after being released from PP2C-mediated inhibition ^42^. However, recent studies suggested that B2 and B3 subgroup RAF-mediated transphosphorylation is essential for the reactivation of ABA-dependent SnRK2.2, SnRK2.3, and SnRK2.6/OST1 ^39–41,43,53^. The high-order mutant of B2 and B3 RAFs, *OK^100^-nonu*, shows an ABA-hyposensitive phenotype in germination and seedling growth, resembling the phenotype of *snrk2.2/2.3/2.6* triple and *pyl112458* high-order mutants ^41^. However, although much insensitive to ABA than the wild type, the stomata of *OK^100^-nonu*, which are still partially closed upon application of ABA, are not identical to those of *snrk2.2/2.3/2.6* triple and *pyl112458* high-order mutants ^41^. Here we showed that the GC-enriched B1 subgroup RAF15 also phosphorylates SnRK2.6/OST1 and mediates ABA signaling in GC. Consistently, the *raf15* single mutant already shows impaired stomatal closure in response to ABA. A very recent study suggested that RAF6, a member of the B3 RAF subgroup, has a dominant role in the rapid vapor pressure difference of GC ^54^. Both *RAF6* and *RAF15* are highly expressed in GCP, compared to MCP (Figure S5A). Thus, the twenty-two B subgroup RAFs and ten SnRK2s may form complex kinase networks, and different cell types may have unique RAF-SnRK2 cascades. The cell-type specialized signaling pathways may ensure that different cells, depending on developmental stage, nutrition status, or microenvironment, respond to similar environmental clues differentially and precisely.

Besides providing clues for discovering the GC-specific RAF15-SnRK2.6/OST1 kinase cascade, our multiomics data also suggest the molecular basis for some long-standing questions in GC structure and function. For example, we found that enzymes in flavone metabolism are highly enriched in GC. Consistently, intermediates in the flavone biosynthesis pathway are also accumulated in GC, which may ensure the maintenance of ROS homeostasis in GC. Our in-depth proteomics and metabolomics data of GC could be a valuable resource for future study of the structure, signaling, function, or even development and differentiation of GC.

## STAR METHODS

Detailed methods are provided in the online version of this paper and include the following:

- KEY RESOURCES TABLE
- RESOURCE AVAILABILITY
  - Lead contact
  - Materials availability
  - Data and code availability
- EXPERIMENTAL MODEL AND SUBJECT DETAILS
  - Plant materials and growth conditions
- METHOD DETAILS
  - Preparation of Arabidopsis GCP and MCP
  - Preparation of Arabidopsis root cell fluorescent protoplasts
  - Preparation of fluorescent protoplasts of rice root cells
  - Protein extraction and digestion
  - LC-MS/MS analysis of proteome
  - Proteomics data analysis
  - Un-targeted metabolomics analysis
  - Stomatal bioassay
  - Protein purification and in vitro kinase assay
  - Protoplast isolation and transactivation assay
- QUANTIFICATION AND STATISTICAL ANALYSIS
  - General statistical analysis
  - Proteomics data analysis.
  - Metabolomics analysis

## SUPPLEMENTAL INFORMATION

Supplemental Figure 1. Identified protein/peptide from GCP by Gdn-Cl and SLS-SDC methods.

Supplemental Figure 2. The enriched GO terms in the proteomes of *DR5 rev::GFP* and *pOs01g0248900:: VENUS-N7* root cells.

Supplemental Figure 3. GO enrichment analysis of proteins enriched in GCP and MCP.

Supplemental Figure 4. Phenylpropane and cutin biosynthesis pathway in MCP and GCP.

Supplemental Figure 5. RAF15 phosphorylate SnRK2.6/OST.

Supplemental Dataset 1. Proteins identified from proteome from 250 to 10,000 GCP by Gdn-Cl and SLS-SDC methods.

Supplemental Dataset 2. Proteins identified from fluorescent protoplasts of *DR5 rev::GFP* root cells.

Supplemental Dataset 3. Proteins identified from fluorescent protoplasts of *pOs01g0248900:: VENUS-N7* root cells.

Supplemental Dataset 4. All peptides identified in GCP.

Supplemental Dataset 5. All peptides identified in MCP.

Supplemental Dataset 6. Quantitative data of proteins identified in GCP and MCP.

Supplemental Dataset 7. GCP enriched proteins identified in this study.

Supplemental Dataset 8. MCP enriched proteins identified in this study.

Supplemental Dataset 9. GO analysis of proteins enriched in GCP and MCP.

Supplemental Dataset 10. The metabolomes of GCP and MCP.

Supplemental Dataset 11. Sequence of primers used in this study.

## ACKNOWLEDGEMENTS

This work was supported by the Strategic Priority Research Program of the Chinese Academy of Sciences, Grant XDB27040106 (to P.W.), and National Natural Science Foundation of China, Grant 31771358 (to P.W.). We are grateful to Drs. Julian Schroeder of University of California, San Diego, for providing the seeds of *pGC1::GFP* transgenic plants, Jiawei Wang of CAS Center for Excellence in Molecular Plant Sciences for providing the seeds of *pOs01g0248900:: VENUS-N7*, Zhaojun Ding of Shandong University for providing the seeds of *WOX5::ERGFP* and *DR5 rev::GFP* transgenic plants. We thank Life Science Editors for editing services.

## AUTHOR CONTRIBUTIONS

P.W. designed the research. H.W., J.R. and Q.G. performed research. H.W., R.L., T.S., C.K., S.D., and C.-P.S. performed data analysis. H.W., R.L., C.K., C.-C.H., and C.-P.S. wrote the manuscript.

## DELARATION OF INTERESTS

The authors declare no competing interests.

## STAR Methods

### KEY RESOURCES TABLE

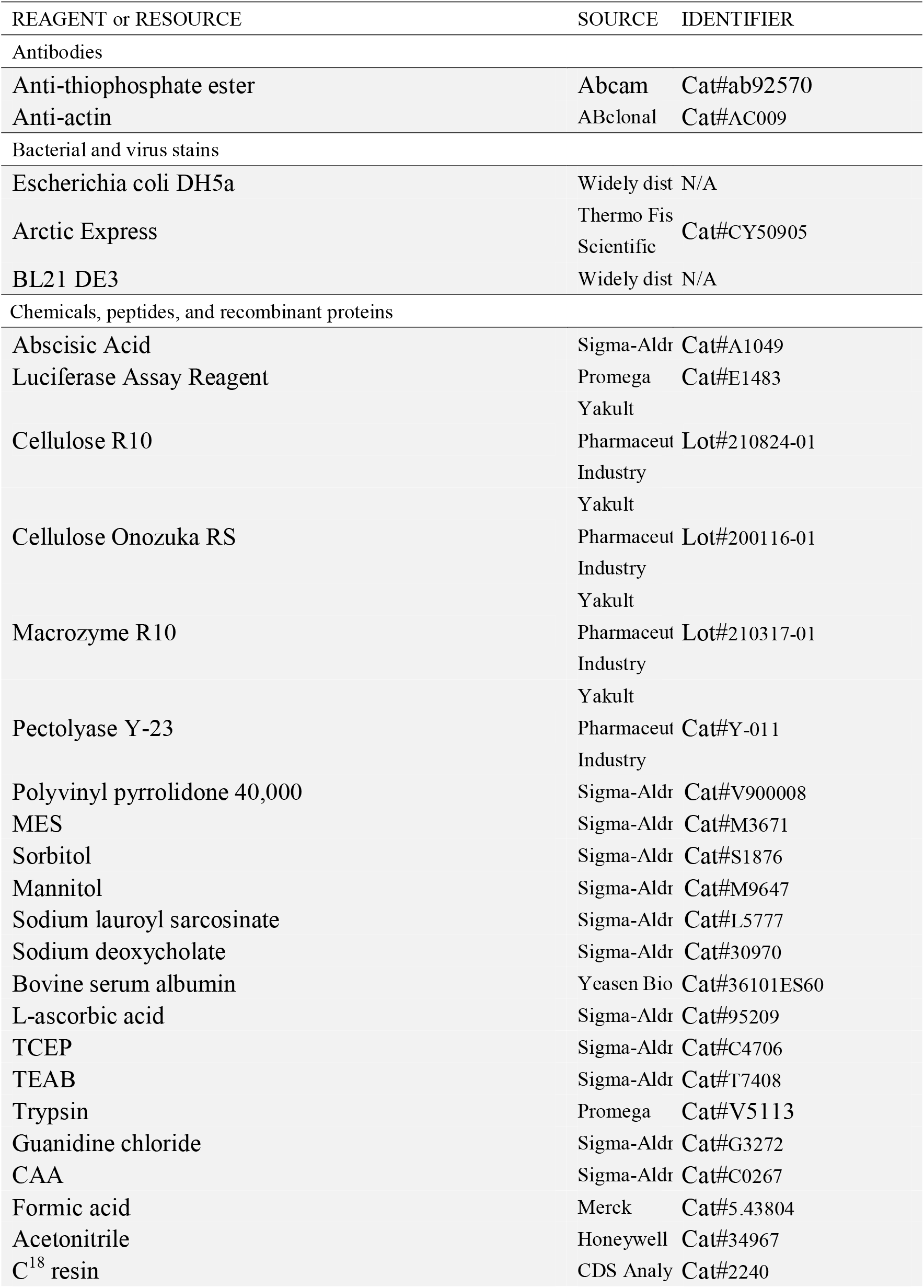
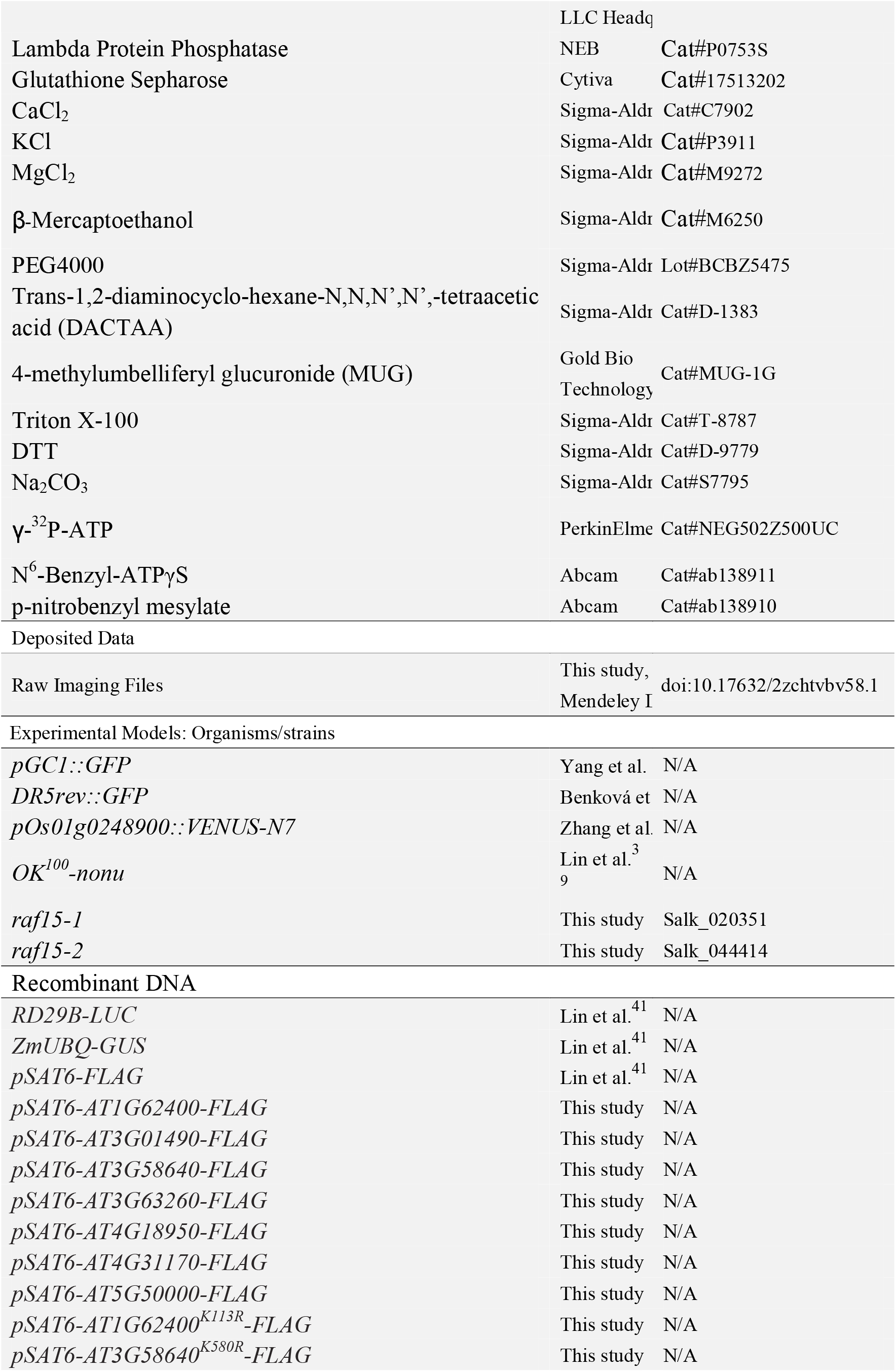
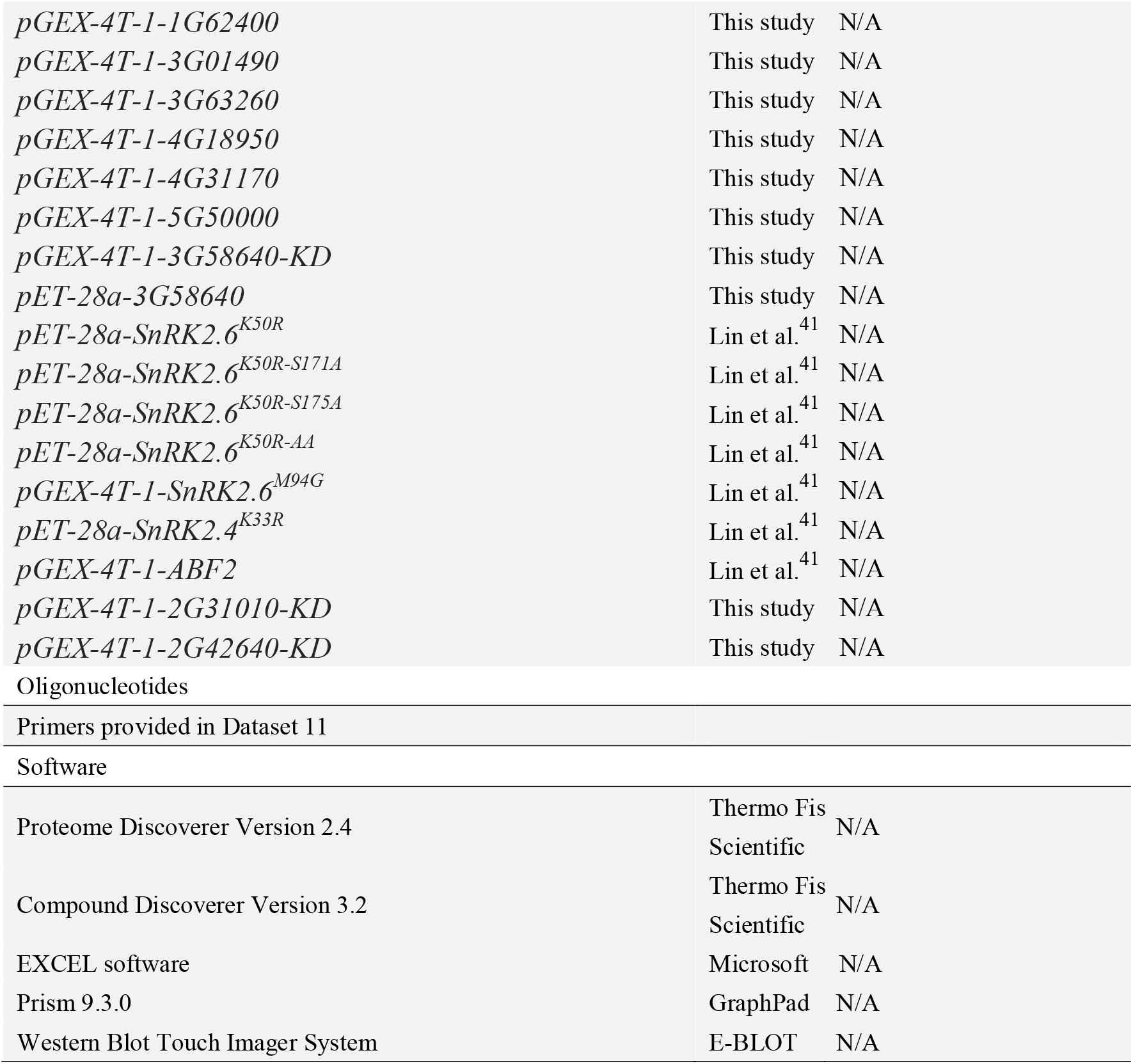

### RESOURCE AVAILABILITY

#### Lead contact

Further information and requests for resources and reagents should be directed to and will be fulfilled by the Lead Contact, Pengcheng Wang.

#### Materials availability

Plasmids and plant materials generated in this study are all available from the Lead Contact upon request.

#### Data and code availability

- The proteomics data were deposited to the ProteomeXchange Consortium with the dataset identifier PXD030489 and PXD030511, the metabolomics data were deposited to MetaboLights with dataset identifier MTBLS3997.
- Any additional information required to reanalyze the data reported in this paper is available from the lead contact upon request.

### EXPERIMENTAl MODEL AND SUBJECT DETAILS

#### Growth conditions and plant materials

The *pGC1::GFP* ^18^ and *DR5rev::GFP* ^21^ seeds were sterilized in 75% ethanol and germinated on half-strength Murashige and Skoog (MS) medium plates with 0.75 agar. After 7 days’ growth on the plates, the *pGC1::GFP* seedlings were transplanted to soil, and rosette leaves were harvested at the indicated time for protoplast isolation.

The seeds of *pOs01g0248900:: VENUS-N7* transgenic rice ^22^ were grown in a half-strength MS liquid medium in the growth chamber at 26 °C with 60% humidity (16 h light/8 h dark). After 7 days, the root tips of the seedlings were harvested for protoplast isolation.

### METHOD DETAILS

#### Preparation of *Arabidopsis* GCP and MCP

The rosette leaves of three-week-old seedlings of *pGC1::GFP* grown in the soil were blended for 10 s five times in 100 mL ice-cold water. The blended mixture was filtered through a 100 μm mesh (Falcon) to remove the broken mesophyll and epidermal cells. The peels were transferred into 20 mL enzyme solution containing 1.35% cellulose R10 (Yakult Pharmaceutical Industry), 0.268% macrozyme R10 (Yakult Pharmaceutical Industry), 0.1% polyvinyl pyrrolidone 40,000 (PVP-40), 0.25% bovine serum albumin (BSA), 0.5 mM L-ascorbic acid, 2.75 mM 2-[N-morpholino] ethanesulfonic acid hydrate-Tris (MES), pH 5.5, 0.275 mM CaCl_2_, 0.275 mM MgCl_2_, 5.5 μM KH_2_PO_4_, and 0.275 M sorbitol at 27 °C with shaking. After 1 h digestion, the peels were filtered with 100 μm mesh and placed into 10 ml enzyme solution II containing 1.3% cellulose Onozuka RS (Yakult Pharmaceutical Industry), 0.0075% Pectolyase Y-23 (Yakult Pharmaceutical Industry), 0.25% BSA, 0.5 mM L-ascorbic acid, 5 mM MES, pH 5.5, 0.5 mM CaCl_2_, 0.5 mM MgCl_2_, 10 μM KH_2_PO_4_ and 0.5 M sorbitol at 27 °C for 2 h. The digestion mixture was filtered with 40 μm mesh (Falcon), and the GCP were collected by centrifugation at 200 × *g* for 5 min. The protoplasts were resuspended in the same solution without enzyme and sorted using cytometry (ARIA III Cell sorter, BD bioscience) with the GFP signal and single-cell pattern to isolate high purity (~ 100%) fluorescent protoplasts. The indicated number of cells were collected by centrifugation and used for protein extraction for LC-MS/MS.

For isolating Arabidopsis MCP, leaf strips were excised from the middle parts of young rosette leaves, dipped in an enzyme solution containing 1.35% cellulase R10 (Yakult Pharmaceutical Industry) and 0.268% macerozyme R10 (Yakult Pharmaceutical Industry), and incubated at room temperature in the dark. The protoplast solution was diluted with an equal volume of W5 solution (2 mM MES, pH 5.7, 154 mM NaCl, 125 mM CaCl_2_, and 5 mM KCl) and filtered through a nylon mesh. The MCP were resuspended with 10 ml W5 buffer and sorted using cytometry (ARIA III Cell sorter, BD bioscience), using chloroplast’s autofluorescence to isolate high purity (~ 100%) fluorescent protoplasts. The indicated number of cells were collected by centrifugation and used for protein extraction for LC-MS/MS.

#### Preparation of Arabidopsis root cell fluorescent protoplasts

To isolate *DR5 rev::GFP* fluorescent protoplasts, the root tip regions (about 0.5 cm from root tip) were cut from the roots of 7-day-old Arabidopsis seedling. The root tips were digested for 2 h in enzyme solution containing 1.5% cellulase R10, 1.5% macerozyme R10, 0.4 M mannitol, 10 mM KCl, 10 mM CaCl_2_ and 0.1% BSA. The protoplasts were filtered with a 40 μm mesh (Falcon) and centrifuged at 500 × *g* for 6 min. The protoplasts were resuspended in 8% mannitol and sorted using cytometry (FACS ARIA III Cell sorter, BD bioscience) with the GFP signal and single-cell pattern to isolate high purity (~ 100%) fluorescent protoplasts. The indicated number of cells were collected by centrifugation and used for protein extraction for LC-MS/MS.

#### Preparation of fluorescent protoplasts of rice root cells

The root tips of 6-day old rice seedlings were harvested and digested for 3 h at room temperature in an enzyme solution containing 4% cellulose Onozuka RS, 1.5% macerozyme R10, 0.4 M mannitol, 0.1 M MES, 10 mM KCl, 10 mM CaCl_2_, and 0.1% BSA with shaking every 30 min. The protoplasts were filtered with a 40 μm mesh (Falcon) and centrifuged at 500 × *g* for 6 min. The protoplasts were resuspended in 8% mannitol and sorted using cytometry (BD bioscience, ARIA III Cell sorter) with the GFP signal and single-cell pattern to isolate high purity (~ 100%) fluorescent protoplasts. The indicated number of cells were collected by centrifugation and used for protein extraction for LC-MS/MS.

#### Protein extraction and digestion

Two sample preparation workflows were compared for the efficiency of protein extraction. GCP and MCP were lysed in SLS-SDC buffer (12 mM sodium lauroyl sarcosinate (SDS), 12 mM sodium deoxycholate (SDC), 100 mM Tris-Cl, pH 8.5) or 6 M guanidine chloride (Gdn-Cl). All the lysis chemicals were dissolved in 100 mM Tris-HCl (pH 8.5). Proteins were reduced and alkylated with 10 mM Tris (2-carboxyethyl) phosphine hydrochloride (TCEP) and 40 mM 2-Chloroacetamide (CAA) at 95 °C for 10 min. Proteins were then subjected to five cycles of ultrasonication pulses of 30 sec and pauses of 30 sec. Protein extracts were 5-fold diluted using 50 mM triethylammonium bicarbonate (TEAB), and trypsin was subsequently added to a final 1:100 (w/w) enzyme-to-protein ratio and incubated at 37 °C overnight. For the Gdn-Cl method, proteins were precipitated using methanol-chloroform precipitation, and the precipitated protein pellet was air dried. The pellet was suspended in SLS-SDC buffer and the SLS-SDC protocol for enzymatic digestion and detergent precipitation followed. For both methods, the digested peptides were desalted using a C18 StageTip.

#### LC-MS/MS analysis of proteome

The peptides were analyzed using Orbitrap Fusion (Thermo Fisher Scientific) or timsTOF pro (Bruker) mass spectrometer. To analyze 1 μg of GCP and MCP digests, the peptides were dissolved in 4 μL of 0.2% formic acid (FA) and injected into an Easy-nLC 1200 (Thermo Fisher Scientific). Peptides were separated on a 15 cm in-house packed column (360 μm OD × 75 μm ID) containing C18 resin (2.2 μm, 100 Å, Michrom Bioresources). The mobile phase buffer consisted of 0.1% FA in ultra-pure water (Buffer A) with an eluting buffer of 0.1% FA in 80% ACN (Buffer B) run over a linear 182 min gradient of 5–35% buffer B at a flow rate of 300 nL/min. The Easy-nLC 1200 was coupled online with an Orbitrap Fusion Tribrid mass spectrometer. The mass spectrometer was operated in the data-dependent acquisition (DDA) mode in which a full-scan MS (from m/z 350–1500 with the resolution of 120,000) was followed by top speed higher-energy collision dissociation (HCD) MS/MS scans of the most abundant ions with dynamic exclusion for 60 s.

For the analysis of the digests of counted 250 to 10,000 GCP cells, the peptides were dissolved in 4 μL of 0.2% FA and injected into nanoflow reversed-phase chromatography, which was performed on an EASY-nLC 1200 system (Thermo Fisher Scientific). Peptides were separated in 120 min at a flow rate of 300 nl min on a 25 cm × 75 μm column with a laser-pulled electrospray emitter packed with 1.6 μm ReproSil-Pur C18-AQ particles. Mobile phases A and B were water with 0.1 % FA and 80:20:0.1 % ACN: water: FA, respectively. The fraction of B was linearly increased from 2% to 27% in 90 min, followed by an increase to 46% in 22 min and a further increase to 100% in 3 min before re-equilibration. Liquid chromatography was coupled online to a hybrid TIMS quadrupole TOF mass spectrometer (timsTOF Pro) via a CaptiveSpray nano-electrospray ion source. The electrospray voltage applied was 1.60 kV. Precursors and fragments were analyzed at the TOF detector, with a MS/MS scan range from 300 to 1500m/z. The timsTOF Pro was operated in parallel accumulation serial fragmentation (PASEF) mode. Precursors with charge states 2 to 3 were selected for fragmentation, and 4 PASEF-MS/MS scans were acquired per cycle. The dynamic exclusion was set to 24 s.

For the analysis of the proteome of fluorescent protoplasts of *DR5 rev::GFP* and *pOs01g0248900:: VENUS-N7* root cells, peptides were separated using a Thermo UltiMate 3000 RSLC nano ProFlow system (Thermo Fisher Scientific) with a gradient comprised an increase from 5% to 38% solvent buffer A (0.1% FA in water) over 80 min, 25%–38% in 15 min, increasing to 99% buffer B (80% ACN, 19.9% H_2_O, 0.1% FA) over 1 min, and then holding at 99% buffer B for the last 9 min, all at a constant flow rate of 100 nL/min. Samples were separated over a 15 cm 100 C18. All samples were analyzed in a nanoscale liquid chromatography (nLC)–MS/MS analysis of 100□min using an Eclipse + FAIMS mass spectrometer (Thermo Fisher Scientific). The mass spectrometer was operated in the data-dependent mode in which a full-scan MS (from m/z 375–1500 with the Orbitrap resolution of 240,000) was followed by Cycle 3 sec higher-energy collision dissociation (HCD) MS/MS scans of the most abundant ions with dynamic exclusion for 60 s.

#### Un-targeted metabolomics analysis

Around 100,000 GCP and 100,000 MCP were suspended in 500 μL of extraction solution (90% methanol). After ultrasonic homogenization for 2 min and vortexing for 10 min, the samples were kept in the dark for 16 h, dried under vacuum, resuspended in 50 μL 40% methanol in water, and centrifuged at 15,000 × *g* for 30 min, 4 °C. The clear supernatants were loaded into injection vials and ready for UHPLC-MS/MS.

For UHPLC-MS/MS assay, the vanquish-flex UHPLC system was coupled to Q Exactive Plus (Thermo Fisher Scientific) for metabolite separation and detection. A Hypersil GOLD column (2.1×100 mm.1.9 um; Thermo Fisher Scientific) was employed for compound separation at 30 °C, and 1 μl of sample was loaded. The mobile phase A was HPLC grade H_2_O with 0.1% (v/v) FA and phase B was HPLC grade ACN. The gradient elution conditions were set as follows: from 0 to 2 min, the mobile phase B increased to 10%; from 2 to 10 min, the mobile phase B increased to 50%; from 10 to 10.1 min the mobile phase B increased to 80%; from 10.1 to 13 min, the mobile phase B was kept at 80%; from 13 to 14 min the mobile phase B increased to 95%; from 14-18 min the mobile phase B decreased to 10%. The flow rate was 0.3 mL/min.

The MS data acquisition was performed by the Q Exactive Plus system. In full scan MS/ddMS^2^ mode, the resolutions of full scan MS and ddMS^2^ were set at 70,000 and 17,500, respectively. The automatic gain control (AGC) target and maximum injection time in full scan MS settings were 1^e6^ and 100 ms, while their values were 2^e5^ and 50 ms in ddMS^2^ settings. The TopN (N, the number of top most abundant ions for fragmentation) was set to 8, and collision energy was set to 20%, 40% and 60%. A heated ESI source was used at positive and negative ion mode. The spray voltage was set as 3.5 KV for positive mode and 3.2 KV for negative mode; the capillary temperature and aux gas heater temperature were set as 320 and 350 °C, respectively. Sheath gas and aux gas flow rate were set at 35 and 15 (in arbitrary units), respectively. The S-lens RF level was 50.

#### Stomatal bioassay

For stomatal aperture assay, rosette leaves of 4-week-old Arabidopsis seedlings were taken. Epidermal strips were peeled out and incubated in buffer containing 50 mM KCl and 10 mM MES, pH 6.15, in a plant growth chamber for 3 h before ABA treatment. Stomatal apertures were measured 2 h after the addition of 5 μM ABA. The apertures of indicated number of stomata per sample were measured by quantifying the pore width of stomata using Image J software. All the experiments were repeated at least three times.

#### Protein purification and *in vitro* kinase assay

For *in vitro* kinase assays, full length coding sequence of SnRK2.6 and kinase domains of RAFs were cloned into either *pGEX-4T-1, pET28a or pET-SUMO vectors* and transformed into BL21 or ArcticExpress cells. The recombinant proteins were expressed and purified as previously described ^39^. For the phosphorylation assay, **recombinant full-length** RAF15, HT1/RAF19, MRK1/RAF48, BHP/RAF27, RAF38, and the kinase domains of RAF13 (aa 492-775) and RAF14 (aa 518-781) were incubated with “kinase-dead” forms of SnRK2.6 with or without Ser to Ala mutations at Ser171 and Ser175 in reaction buffer (25 mM Tris HCl, pH 7.4, 12 mM MgCl_2_, 2 mM DTT), with 1 μM ATP plus 1 μCi of [γ-^32^P] ATP for 30 min at 30°C. Reactions were stopped by boiling in SDS sample buffer and proteins were separated by 10% SDS-PAGE. Primers used in RAF construction are listed in Supplementary Dataset 11.

For the dephosphorylation assay, SnRK2.6^M94G^ coated on Glutathione Sepharose (Cytiva) were dephosphorylated with Lambda Protein Phosphatase (λPP) for 30 min and the λPP was removed by washing three times with protein buffer (25 mM Tris HCl, pH 7.4, 150 mM NaCl). To detect the effects of RAF15 on SnRK2.6^M94G^ thiophosphorylation and activity, recombinant GST-RAF15-KD (aa 548-809) was incubated with pre-dephosphorylated SnRK2.6^M94G^ for 30 min in reaction buffer (25 mM Tris HCl, pH 7.4, 12 mM MgCl_2_, 2mM MnCl_2_, 0.5 mM DTT, 50 μM ATP, 50 μM N^6^-Benzyl-ATPγS). Then ABF2 was added to the reaction and incubated for an additional 30 min. This phosphorylation reaction was stopped by adding EDTA to a final concentration of 25 μM. A final concentration of 2.5 mM p-nitrobenzyl mesylate (Abcam, ab138910) was added to proceed the alkylating reaction for 1 h at room temperature. Samples with SDS sample buffer were boiled and separated by SDS-PAGE, transferred to Polyvinylidene fluoride (PVDF) membrane, and immunoblotted with antibodies against thiophosphate ester (Abcam, ab92570).

#### Protoplast isolation and transactivation assay

Protoplast isolation and transactivation assays were performed as previously described ^19^. Briefly, protoplasts were isolated from leaves of 4-week-old plants grown under a short photoperiod (10 h light at 23°C/14 h dark at 20°C). Leaf strips were excised from the middle parts of young rosette leaves, dipped in an enzyme solution containing cellulase R10 (Yakult Pharmaceutical Industry) and macerozyme R10 (Yakult Pharmaceutical Industry), and incubated at room temperature in the dark. The protoplast solution was diluted with an equal volume of W5 solution (2 mM MES, pH 5.7, 154 mM NaCl, 125 mM CaCl_2_, and 5 mM KCl) and filtered through a nylon mesh. The flow-through was centrifuged at 100 *g* for 2 min to pellet the protoplasts. Protoplasts were resuspended in W5 solution and incubated for 30 min. 100 μL of protoplasts suspended in MMG solution (4 mM MES, pH 5.7, 0.4 M mannitol, and 15 mM MgCl_2_) were mixed with the plasmid mix and added to 110 μL PEG solution (40% w/v PEG-4000, 0.2 M mannitol, and 100 mM CaCl_2_). The transfection mixture was mixed completely by gently tapping the tube followed by incubation at room temperature for 5 min. The protoplasts were washed twice with 1 mL W5 solution. The *RD29B-LUC* (7 μg of plasmid per transfection) and *ZmUBQ-GUS* (1 μg per transfection) were used as an ABA-responsive reporter gene and as an internal control, respectively. For wild type and mutated *RAF, SnRK2.6* plasmids, 3 μg per transfection were used. After transfection, protoplasts were incubated for 5 h under light in washing and incubation solution (0.5 M mannitol, 20 mM KCl, 4 mM MES, pH 5.7) with or without 5 μM ABA. The mutations were introduced into wild type RAFs using the primers listed in Supplementary Dataset 11.

#### Primer list

Dataset S11 presents the sequences of the primers used in this study.

### QUANTIFICATION AND STATISTICAL ANALYSIS

#### General statistical analysis

Statistic significance of relative luciferase activity, phosphatase activity, fresh weight, Rosetta diameter, relative intensity, ion leakage, gene expression was examined by Student’s t test (*p < 0.05, **p < 0.01, ***p < 0.001).

#### Proteomics data analysis

The raw files were searched directly against the Arabidopsis thaliana database (TAIR10 with 35,386 entries) with no redundant entries using SEQUEST HT search algorithm on Proteome Discoverer software (version 2.4). Peptide precursor mass tolerance was set at 10 ppm, and MS/MS tolerance was set at 0.02 Da. Three missed cleavage sites of trypsin were allowed. Search criteria included a static carbamidomethylation of cysteine (+57.0214 Da) and variable modifications of oxidation (+15.9949 Da) on methionine and phosphorylation (+79.9663 Da) on serine, threonine, and tyrosine. The false discovery rates (FDR) of proteins and peptides were set at 1% FDR.

#### Metabolomics analysis

Following LC-MS analysis, raw data were collected and processed using Compound Discoverer 3.2 (Thermo Fisher Scientific) with the metabolite databases including mzCloud, mzVault, Masslist, and Chemspider. Principal component analysis (PCA), heatmap analysis of metabolites, and volcano plots were performed using R (version 4.0) software.

