## Supplementary material for "A cell type-specific multiomics uncovers a guard cell-specific RAF15-SnRK2.6/OST1 kinase cascade": Supplmental Figure 1-5

**Ssupplemental Information**

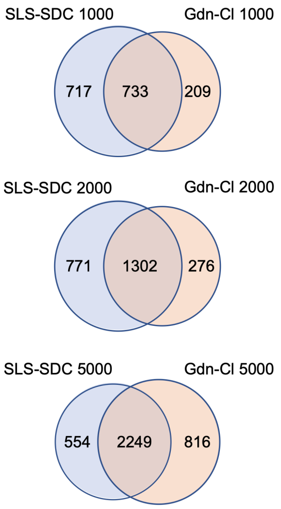


**Supplemental Figure 1**. **Venn diagram showing the overlap between the proteins identified from the indicated number of fluorescent protoplasts using the SLS-SDC and Gdn-Cl methods.**

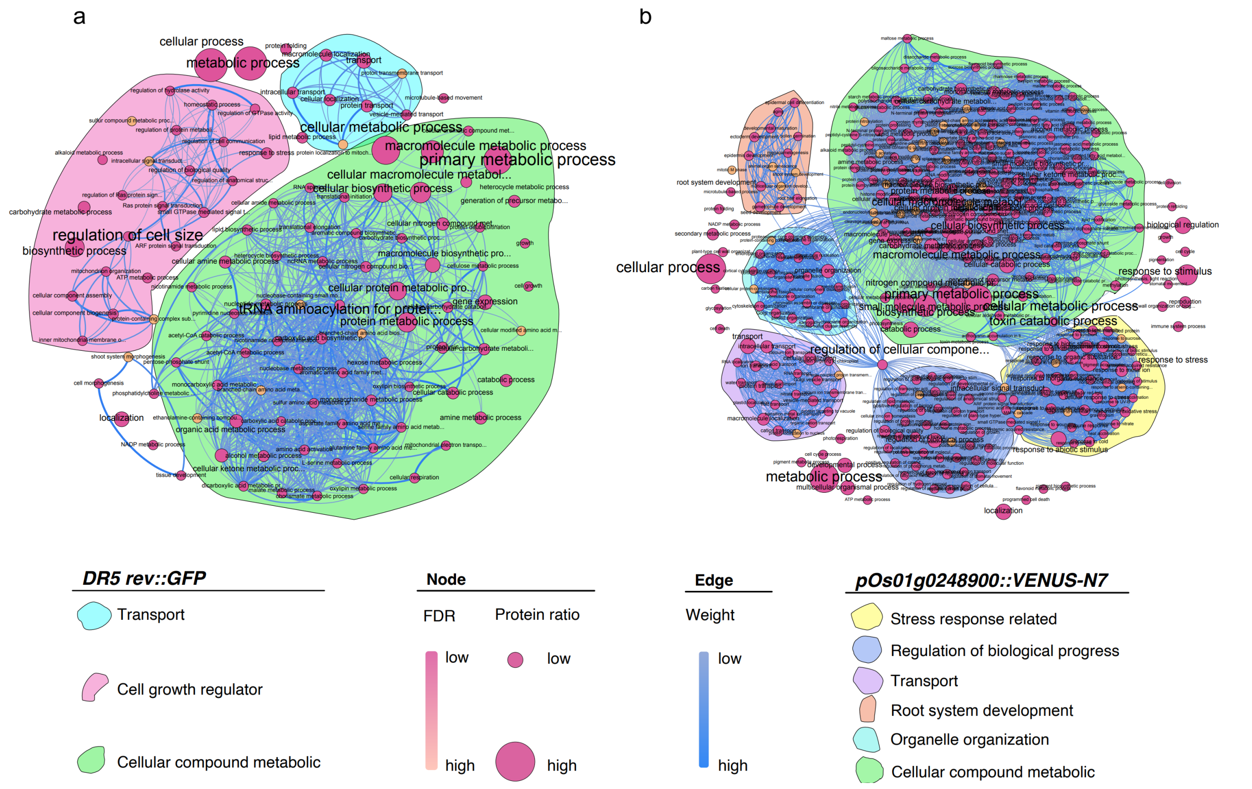


**Supplemental Figure 2. GO analysis of proteomes of *DR5 rev::GFP* and *pOs01g0248900::VENUS-N7* root cells.**

**A.** GO analysis of proteome in fluorescent protoplasts of *DR5 rev::GFP* transgenic plants. **B.** GO analysis of proteome in fluorescent protoplasts of *pOs01g0248900::VENUS-N7* transgenic plants. The original GO enrichment results are filtered by REViGO to remove semantic redundancy. The circular dots represent GO terms, and the lines between the dots represent the connection between GO terms. The bubble size is determined by the significance of enrichment (number of proteins in corresponding GO term/total number of input). The bubble color is decided by FDR (BH-method). Independent networks or sub-networks are manually circled with irregular graphics.

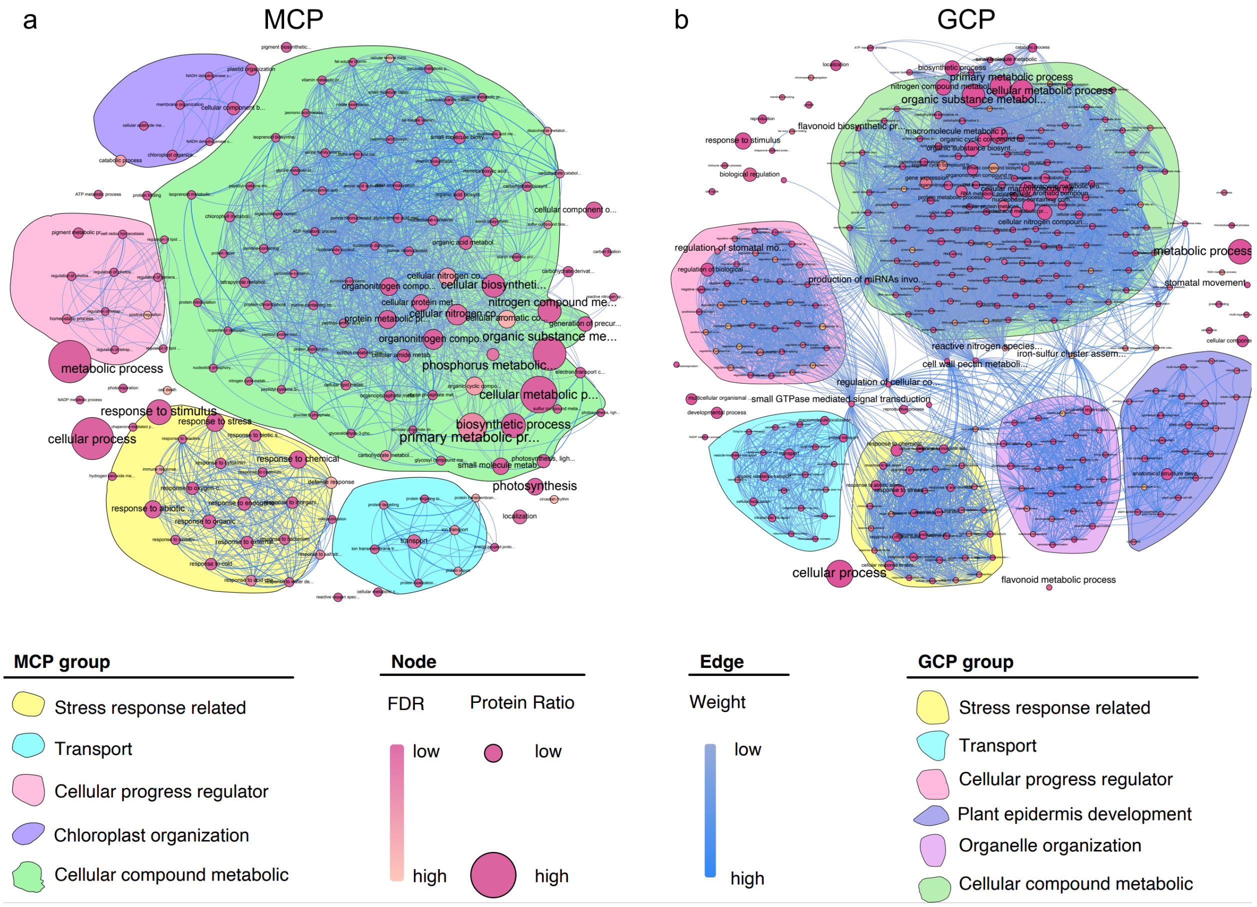


**Supplemental Figure 3. GO enrichment analysis of proteins enriched in GCP and MCP. A.** GO enrichment analysis of proteins enriched in GCP. **B.** GO enrichment analysis of proteins enriched in MCP. The original GO enrichment results are filtered by REViGO [1] to remove semantic redundancy. The circular dots represent GO terms, and the lines between the dots represent the connection between GO terms. The bubble size is determined by the significance of enrichment (number of proteins in corresponding GO term/total number of input). The bubble color is decided by FDR (BH-method). Independent networks or sub-networks are manually circled with irregular graphics.

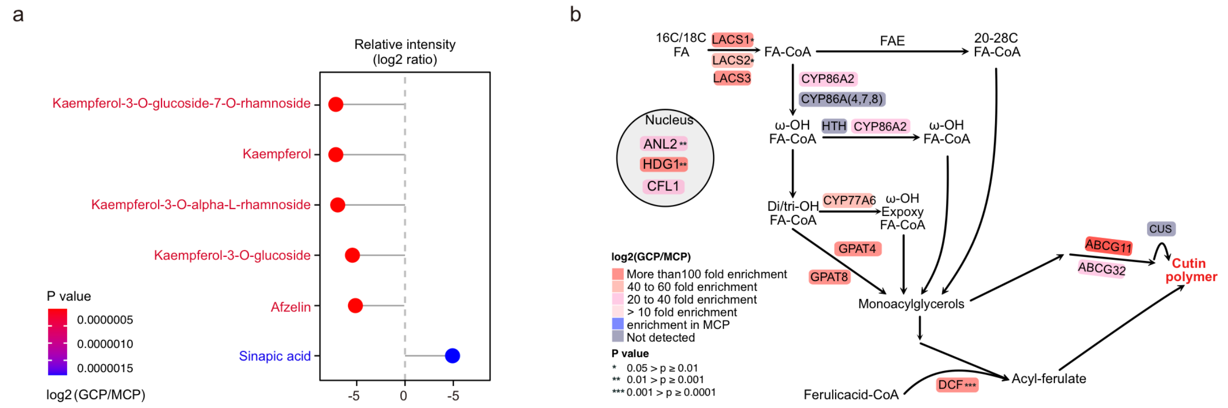


**Supplemental Figure 4. Phenylpropane and cutin biosynthesis pathway in MCP and GCP.** **A.** The amount of phenylpropane compounds detected by non-target metabolites in GCP and MCP. The x-axis represents the relative abundance (log2 ratio), and the color is determined by the *p* value (unpaired two-tailed *t*-test). **B.** The protein-metabolite joint pathway analysis of cutin biosynthesis between GCP and MCP. The metabolites, proteins, and their relationships in the figure on the left are derived from the description of Fich et al., 2016.

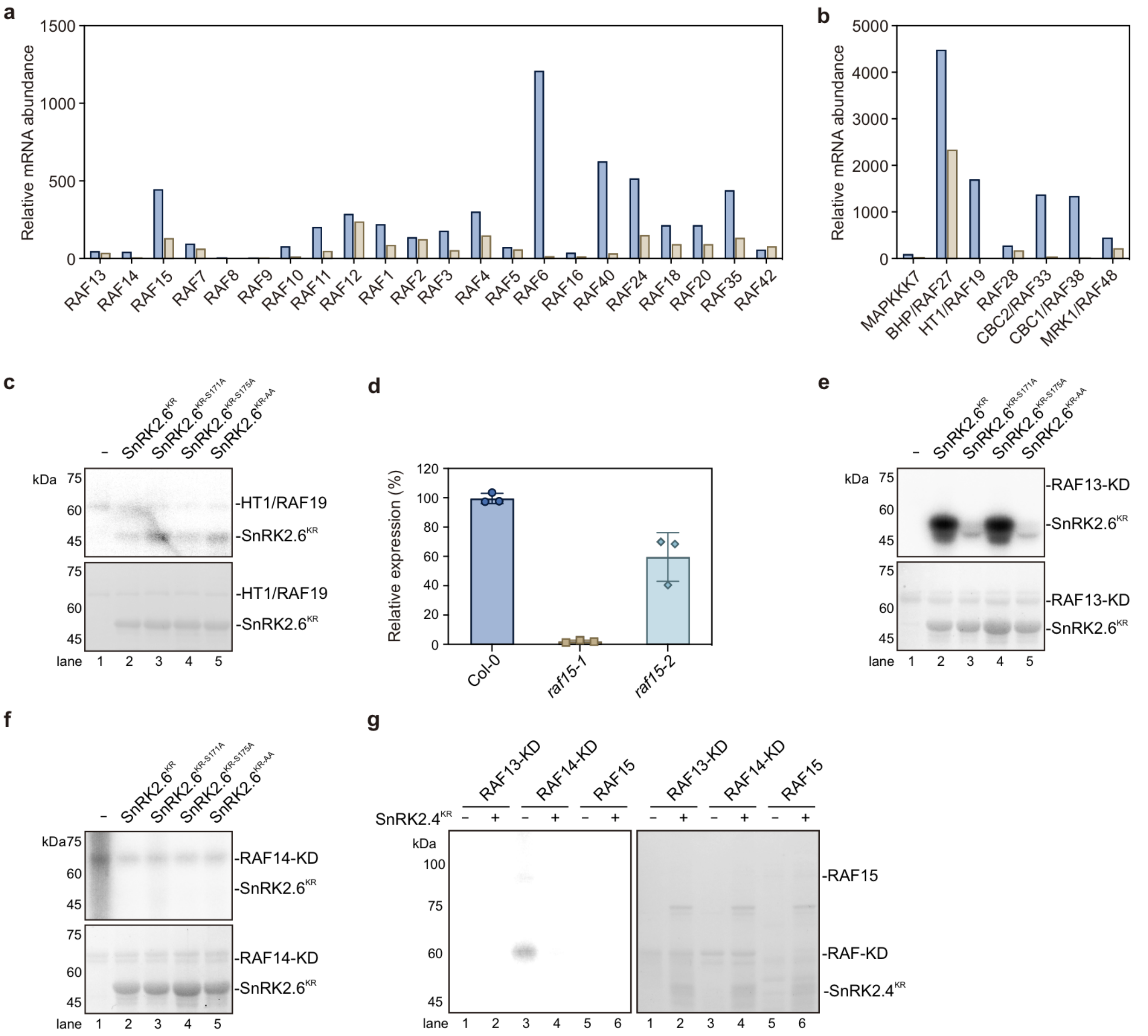


**Supplemental Figure 5. RAF15 and RAF13 phosphorylate SnRK2.6/OST1 in vitro.**

**A.** The relative abundance of B1, B2, B3, and B4 *RAF* transcripts in GC and MC. **B.** The relative abundance of A and C subgroup *RAF* transcripts in GC and MC. The expression of *RAFs* shown in **A.** and **B.** was obtained from the Arabidopsis eFP browser web site http://bar.utoronto.ca/efp/cgi-bin/efpWeb.cgi. **C.** Recombinant GST-HT1/RAF19-KD were used to phosphorylate wild type and mutated SnRK2.6$^{KR}$ in the presence of [$\gamma$-$^{32}$p]ATP. Autoradiograph (upper) and Coomassie staining (bottom) show phosphorylation and loading, respectively, of purified GST-HT1/RAF19-KD and HIS-SnRK2.6$^{KR}$. **D.** Expression of *RAF15* in *raf15-1* and *raf15-2* mutants. **E.** Recombinant GST-RAF13-KD were used to phosphorylate wild type and mutated SnRK2.6$^{KR}$ in the presence of [$\gamma$-$^{32}$p]ATP. Autoradiograph (upper) and Coomassie staining (bottom) show phosphorylation and loading, respectively, of purified GST-RAF13-KD and HIS-SnRK2.6$^{KR}$. **F.** Recombinant GST-RAF14-KD were used to phosphorylate wild type and mutated SnRK2.6$^{KR}$ in the presence of [$\gamma$-$^{32}$p]ATP. Autoradiograph (upper) and Coomassie staining (bottom) show phosphorylation and loading, respectively, of purified GST-RAF14-KD and HIS-SnRK2.6$^{KR}$. **G.** Recombinant GST-RAF13-KD, GST-RAF14-KD, and HIS-RAF15, were used to phosphorylate wild type and mutated SnRK2.4$^{KR}$ in the presence of [$\gamma$-$^{32}$p]ATP.
